# Rising from the depths: recent abundance and distribution trends of the bluntnose sixgill shark (*Hexanchus griseus*) in the Mediterranean Sea

**DOI:** 10.1101/2025.07.04.663171

**Authors:** Iris L. Jia, Francesco Ferretti

## Abstract

Sharks play a crucial role in the structure of marine ecosystems but are heavily threatened by overfishing globally. Due to their slow life history traits, even low fishing pressure can have an extreme influence on shark populations. The Mediterranean Sea, which has had historical overexploitation over the course of many centuries, is considered a hotspot for elasmobranchs. One particular species is the bluntnose sixgill shark, *Hexanchus griseus*, which has recently become more prominent in sightings and landings. Because of its deep range, *H. griseus* is difficult to study *in situ* so we used opportunistic data from sharkPulse, a crowd-sourcing platform that collects and organizes image-based shark occurrence records from all over the world, to better understand the abundance and distribution of *H. griseus* over the past decade. Additionally, we explored historical data to create a baseline of the species’ abundance starting from the late 1800s. We found that *H. griseus* underwent a quadratic-shaped change in their abundance where they started out as a rare catch, became increasingly more prominent in fisheries data, and now is declining like many other cartilaginous fishes in the Mediterranean Sea. This work is important because *H. griseus* is not yet threatened according to the IUCN so research is not heavily focused on this species but the decreasing trend we detected here is concerning while in line with the degree of exploitation of one of the most exploited regions of the world, as well as illustrative of a broader ecological process that will likely occur in many other large marine ecosystems worldwide.

## INTRODUCTION

Despite playing a crucial role in the function and structure of many marine ecosystems, sharks and their relatives are facing threats unprecedented in their 420-million-year history (Nuez *et al.* 2023). One of the most severe threats is overfishing, which is driving over 1/3 of Chondrichthyian species to global extinction (Dulvy *et al.* 2021). Due to their slow life history strategies, characterized by late sexual maturity, low fecundity and long gestation, cartilaginous fish are affected by low fishing pressure (Bargnesi *et al.* 2019). As top- and meso-predators in many different marine ecosystems, the loss of shark populations can lead to severe ecosystem consequences across multiple trophic levels (Heithaus *et al.* 2008). Shark overexploitation was linked to the collapse of a century-old bay scallop fishery in the northwestern Atlantic (Myers *et al.* 2007). The loss of coral due to mesopredator release and subsequent decrease of herbivores occurred after the removal of sharks in northwest Australia (Ruppert *et al.* 2013), and many other food web disturbances globally were seen in correspondence with shark population depletions (Ferretti *et al.* 2010). Although there is evidence of far-reaching ecological consequences following the removal of sharks, their conservation status remains poor with 32.6% of species threatened with extinction (IUCN; Dulvy *et al*. 2021) and the appropriate management strategies are still debated (Birkmanis *et al.* 2020).

One key hotspot for elasmobranchs is the Mediterranean Sea (Mancusi *et al.* 2023). Although the Mediterranean Sea comprises less than 1% of the global ocean, 47 species of sharks have been recorded here (Bargnesi *et al.* 2019). Unfortunately, in this region, sharks and rays are heavily impacted by numerous threats, including but not limited to habitat loss, eutrophication, climate change, and invasive species (Nuez *et al.* 2023). Additionally, the Mediterranean Sea has one of the longest human exploitation histories in the world (Colloca *et al.* 2017). The past centuries have shown steep declines of shark populations within the Mediterranean (Bargnesi *et al.* 2022, Ferretti *et al.* 2008, 2013), particularly due to an increasing and uncontrolled fishing pressure, which has resulted in the Mediterranean having the highest proportion of threatened species (Nuez *et al.* 2021).

One species that emerged more prominently in recent years is the bluntnose sixgill shark (*Hexanchus griseus)*, which is a large, deep-benthic shark with a worldwide geographic distribution (Griffing *et al.* 2019). *H. griseus* is currently categorized on the International Union for Conservation of Nature (IUCN) Red List of Threatened Species as Near Threatened (Finucci *et al*. 2020) and regionally listed as Least Concerned in the Mediterranean assessment (Soldo *et al.* 2016). Given its ability to feed on diverse prey (Rodríguez-Cabello *et al*. 2018), it is thought to be a generalist apex predator and able to shape ecosystems in its spatial niche. This species is known to inhabit depths ranging from the surface to 2500 m with a typically observed range of 180-1100 m (Nakamura *et al.* 2015). They are also shown to undergo diel vertical migration where they descend to depths below 500 m during the day and ascend to 200-350 m during the night (Coffey *et al.* 2020). This migration exposes them to a wide range of temperatures and dissolved oxygen conditions (Coffey *et al.* 2020) and puts them at risk of overlapping with fisheries. Since the 1970s, with declines of shallow-water stocks, fishery operations have increased depth (Bottaro *et al.* 2023). Although most Mediterranean fisheries, which can reach depths of 800-1000 m (Farrag 2022), do not directly target elasmobranchs, many species are incidentally captured (Albonetti *et al.* 2023). *H. griseus* in particular is a common bycatch species of bottom and mid-water trawlers but the species has also been caught by gillnets, trammel nets, longlines, handlines and traps (Nuez *et al.* 2023). Even though *H. griseus* is currently assessed as Least Concerned in the Mediterranean, the potential of being caught in such a wide range of fishing gear with little to no international protection warrants the attention of conservation efforts. Neglecting non-threatened species can lead to severe consequences of their actual conservation status, an example being the blue shark which went from Vulnerable in 2006 to Near Threatened in 2009 to Critically Endangered in 2016 (Nuez *et al.* 2023).

With the depletion of other large predatory sharks in the Mediterranean Sea in the past 50-200 years as a result of intense overfishing (Ferretti *et al.* 2008), there exists an opportunity for deep-sea species like *H.griseus* to expand its ecological niche. Records from a survey study conducted from 2000 to 2009 showed that *H. griseus* was one of the most common species landed/seen in the central Mediterranean Sea (Sperone *et al.* 2012). A similar phenomenon was observed in South Africa where the broadnose sevengill shark (*Notorynchus cepedianus*), which is a similar ecotype to *H. griseus*, became increasingly abundant following long periods of white shark disappearance (Hammerschlag *et al.* 2019). Although Hammerschlag *et al.* worked on a much smaller temporal and spatial scale, their results suggest mechanisms that can scale up in space and time.

To better understand the abundance and distribution of *H. griseus* with relation to other large predatory sharks and fishing pressure, we used sharkPulse, a crowdsourcing platform collecting and organizing shark photo sightings from around the world (Bargnesi *et al.* 2022, sharkPulse.org). We analyzed the Mediterranean portion of the database to estimate spatiotemporal changes in abundance and distribution of *H. griseus* in the region and compared these trends with historical population baselines extracted from old published literature. We expected to see a quadratic change in *H. griseus* abundance, initially increasing due to the long exploitation history of the Mediterranean Sea which has strongly impacted other large predatory sharks (important predators and competitors) but lightly affected the *H. griseus* due to its deep water distribution, and eventually declining when Mediterranean fisheries expanded and intensified.

## METHODS

We initially extracted all shark occurrence records from the Mediterranean Sea contained in the sharkPulse database. SharkPulse is a crowdsourcing platform sourcing digital pictures from online repositories and transforming them into shark occurrence records (Ferretti *et al.* in prep); the species identification of these records are validated by members of the sharkPulse team and elasmobranch researchers. SharkPulse data are stored in a PostgreSQL relational database (version 9.5.19). Valid sharkPulse records used in this study required the date, latitude, longitude, image, and source (i.e. submission through app, from a website, Instagram, etc.). Additionally, we downloaded sightings from the Global Diversity Information Facility (GBIF) and the IUCN. The IUCN shapefile gave us an expectation of where *H. griseus* was likely to occur. Plotting the GBIF data was useful in understanding the global patterns in the distribution of *H. griseus* and whether what we were seeing with sharkPulse was illustrative of actual population dynamics or sampling and observation efforts.

Due to the slow life history traits of *H. griseus* like a two-year reproductive cycle (Santander-Neto *et al.* 2023) and age of maturity being 11-14 years for males and 18-35 for females, population changes due to any number of external influences are expected to unfold over many decades. Our analysis of sharkPulse records looks at a brief snapshot of recent data to better understand long-term population trends. We extracted information regarding the length, weight, age, and sex of individuals from the records’ source websites when available. Missing lengths when weights were available and vice versa were additionally predicted using a length-weight relationship (W = aL^b) fitted to a pool of 45 individuals that contained both length and weight measurement. Due to only 13 entries in regards to age, we used the von Bertalanffy growth curve to estimate age for all sharkPulse entries that contained length. The coefficients for the von Bertalanffy growth curve were taken from FishBase and this allowed us to divide our records into mature and immature individuals using a length at maturity of 4.51 m and 3.10 m for females and males respectively (COSEWIC 2007). To better understand where our data were coming from, we divided the records into two umbrella categories: catch records and sightings. Catches were further categorized as records from a fish market and in the process of being fished either onboard a boat or landed at a port. Sightings were divided into strandings or dive-related (shark was seen underwater).

We used the opportunistic data sourced from sharkPulse to estimate temporal and spatial trends of *H. griseus* abundance. This analysis was limited from 2012 to 2021 due to limited data prior to sharkPulse’s activation. A lower observation effort was also expected during 2019-2021 due to the COVID-19 pandemic. We fitted a generalized linear model (GLM) with year as a factor and a GLM with year as a continuous variable to the number of *H. griseus* observations as a function of latitude, longitude, depth and year. The annual number of shark sightings recorded in sharkPulse was used as a proxy for observation effort. This approach, which is similar to the one used in Bargnesi *et al.* (2022), assumes that the detection rate of *H. griseus* from the opportunistic data would increase with an increasing amount of shark observations in general. Both models were fitted using a Poisson distribution with a logarithm link function. Furthermore, we fitted the data with a generalized additive model (GAM) having the same response variable and covariates. However, this time, latitude and longitude were included as interactive splines in an effort to predict the distribution of *H. griseus* populations across the Mediterranean Sea over the period 2012 - 2021.

We supplemented our sharkPulse analysis with historical information about the perceived abundance of *H. griseus* as described by early naturalists and scientists working in the Mediterranean Sea by compiling literature from different regions in the Mediterranean starting from 1883 and ending in 2012. These historical data sources were mainly papers and books published on the fishing trends in the Mediterranean. We tabulated the researchers’ perception of the abundance of *H. griseus* over the course of nearly two centuries (Table 1) by ranking the researchers’ perceived abundances with a four-level class coding system (very rare, rare, common, and very common) that was used in Fortibuoni *et al.* (2010). Then we assigned numeric values to each class (very rare = 0.2, rare = 0.4, common = 0.6, very common = 0.8) in order to find the average perception over 50-year periods. Additionally, we recorded the gear type, region, and study period of each reference when available. This was useful to contextualize the perceived abundance into the fishing conditions generating encounters with *H. griseus*, understand what regions were seeing more or less of the shark, and whether their populations were more or less prominent compared to modern times. Statistical analysis, mapping, and plotting were all conducted using R (RStudio, Version 2023.06.2+561).

**Table 1.** Data from 1883-2012 regarding the perceived abundance of *H. griseus* in various regions of the Mediterranean. The perceived abundance was categorized into four levels: very rare, rare, common, very common. (* for UNEP Regions signify that *H. griseus* was mentioned in those countries’ reports)

| Reference | Perceived Abundance | Region | Gear Type | Study Period (if applicable) |
| --- | --- | --- | --- | --- |
| Faber (1883) | rare | Adriatic Sea |  |  |
| Brusina (1888) | rare | Croatia |  |  |
| Parona (1898) | very rare | Liguria |  |  |
| Paolucci (1901) | very rare | Italian Area of Adriatic Sea |  |  |
| Ninni (1912) | rare | Adriatic Sea |  |  |
| Pastrovic (1913) | not mentioned | Adriatic Sea |  |  |
| Dancona and Razzauti (1939) | not mentioned | Tyrrhenian Sea | trawl |  |
| Dieuzeide *et al.* (1953) | rare | Algerian Coasts | trawl and longline | 1923 - 30 years prior to publication |
| Kirincic and Lepetic (1955) | very rare | Southern Adriatic Sea | longline |  |
| Capape (1989) | rare | throughout entire Mediterranean Sea |  |  |
| Fergusson and Marks (1996) | common | Maltese Archipelago (Central Mediterranean) | deepwater trawl and bottom longline | 1983-1992 |
| Bello (1999) | rare | Southwestern Adriatic Sea | trawl | literature started from 1975 (technically from 1948 as Šoljan's work was an updated edition of the work done in 1948) |
| Barrull *et al.* (1999) | common | Catalan Waters | trawl and longline | 1987-2000 |
| Capape *et al.* (2000) | rare | Greece, Italy, Spains | trawl | 1998-1999 |
| Megalofonou *et al.* (2000) | rare | Languedoc coast (Southern France) | longline | literature starting from 1860; observations starting from 1988 |
| Machias *et al.* (2001) | rare | Northeastern Mediterranean |  | 1995-1998 |
| UNEP (2002) | rare | Albania, Croatia*, Greece, Cyprus, Lebanon, Morocco*, Tunisia | trawl |  |
| Megalofonou *et al.* (2005) | very rare | 9 sampling areas across the Mediterranean |  | 1998-1999 |
| Megalofonou *et al.* (2005) | not mentioned | Eastern Mediterranean | drifting longline | 1998-2001 |
| Hadjichristophorou (2006) | common | North and West coasts of Cyprus |  | 1980-2004 |
| Bradaï *et al.* (2006) | common | Gulf of Gabès | bottom trawl, gillnet and longline | 1995-2004 |
| Saad *et al.* (2006) | very common | Syrian coast of Mediterranean |  | 2001-2004 |
| Golani (2006) | common | Israel coasts |  |  |
| Schembri (2006) | rare | Maltese Islands (Central Mediterranean) |  |  |

## RESULTS

A total of 242 *H. griseus* occurrence records reporting latitude and longitude coordinates and information of observation time and date were extracted from the sharkPulse database. The coordinates ranged between 30 to 46 °N and 6 °W to 36 °E. Three observations were excluded from this pool as they were recorded before 2012. The remaining 239 entries were summarized in Table 2 aggregated by country of origin, observation type and year (Table 2). Most of the observations were from Greece, followed by Turkey and Italy. All countries had a higher number of catch observations compared to sightings; the total number of catch observations were seven times greater than sightings (Table 2, Figure 2). The sharkPulse data is highly concentrated in the Mediterranean Sea while the GBIF data is more focused in southern Africa as well as the west coast of the United States and South America (Figure 1). These are areas with steep continental slopes, which makes the habitat most frequently used by the species more proximal to fisheries domains and the reach of human ocean use. The IUCN distribution also shows that *H. griseus* has been seen in Australian and east Asian waters but data from sharkPulse and GBIF are lacking in these areas (Figure 1).

**Figure 1.**
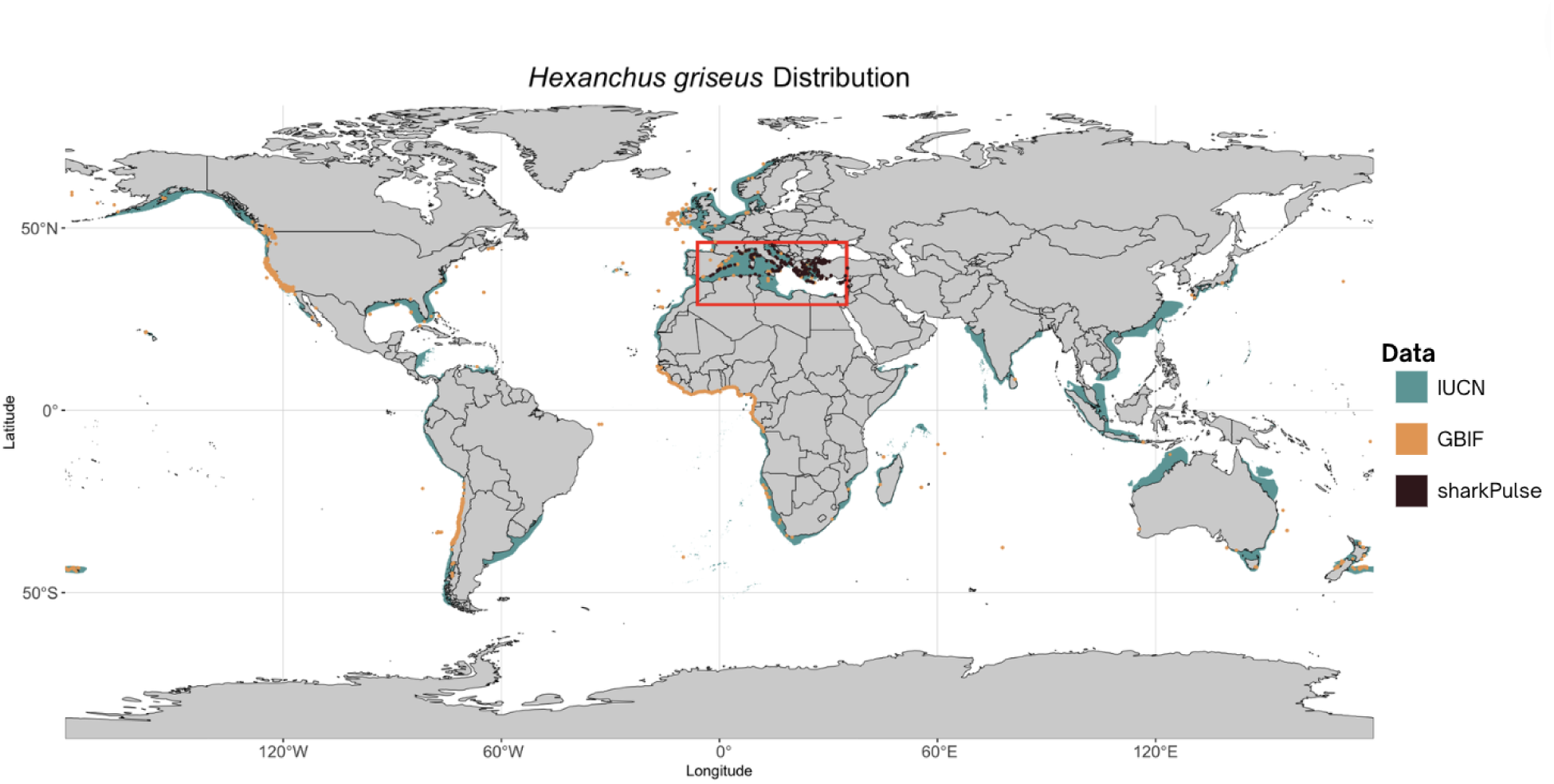
Map of the current IUCN distribution range of *Hexanchus griseus* with an overlay of all sharkPulse and GBIF records. The red box highlights the Mediterranean Sea, our study region.

**Figure 2.**
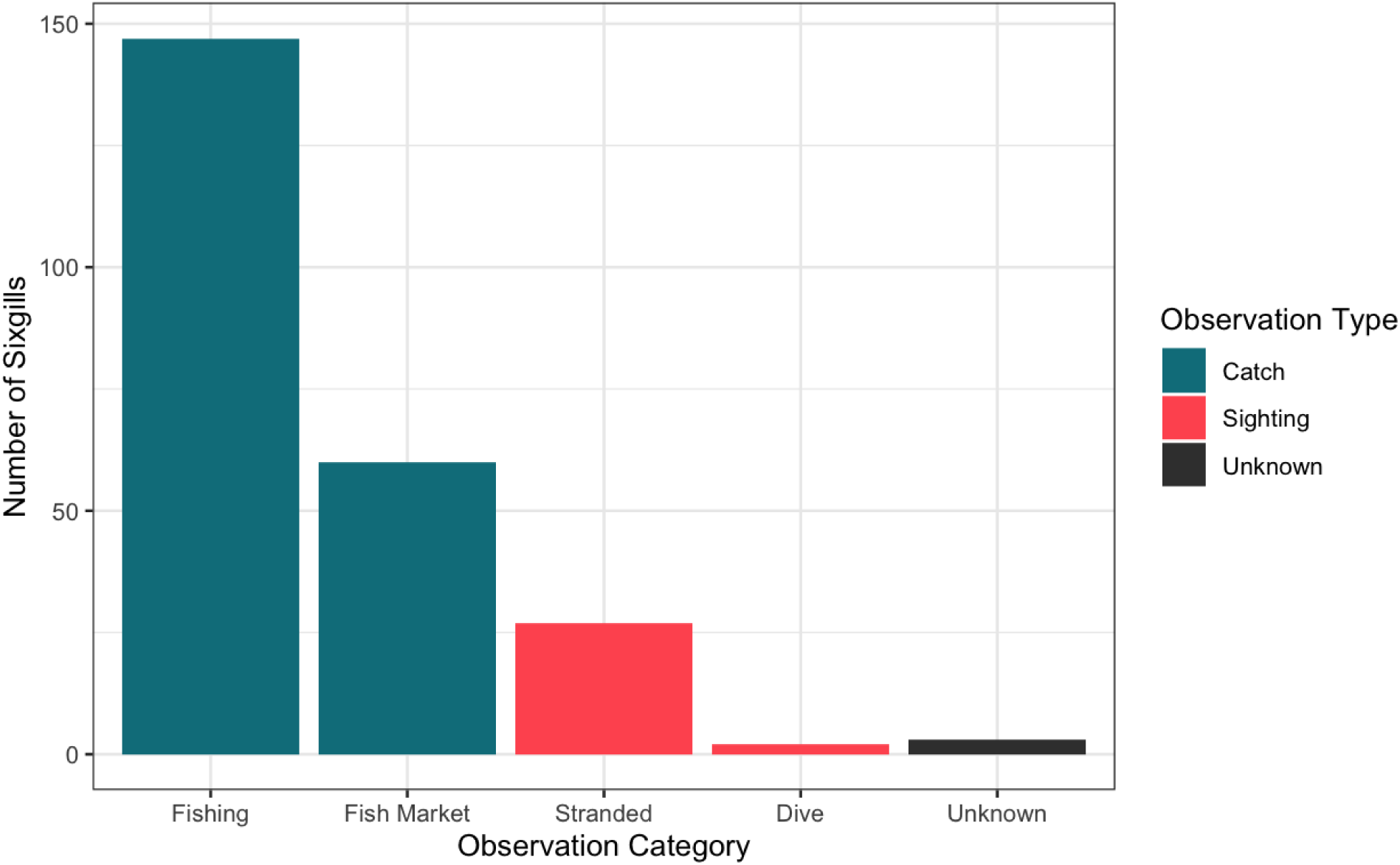
Bar graph of the additional categories under the observation types of catch and sighting.

**Table 2.** Summary table of sharkPulse *Hexanchus griseus* observations and their country of origin; the entries were separated into catch and sighting categories and also broken down by year. Three of the 239 entries were excluded because their observation type was unknown. The records were from Turkey in 2014, Turkey in 2021, and Greece in 2021.

| Country of Origin | Observation Type |  | Year |  |  |  |  |  |  |  |  |  |  |  |
| --- | --- | --- | --- | --- | --- | --- | --- | --- | --- | --- | --- | --- | --- | --- |
|  | Catch | Sighting | 2012 | 2013 | 2014 | 2015 | 2016 | 2017 | 2018 | 2019 | 2020 | 2021 | 2022 | 2023 |
| Greece | 64 | 6 | 4 | 2 | 7 | 1 | 3 | 20 | 8 | 16 | 4 | 2 |  | 3 |
| Turkey | 52 | 1 | 12 | 9 | 5 | 1 | 1 | 6 | 15 | 4 |  |  |  |  |
| Italy | 36 | 15 | 4 | 12 | 4 | 1 | 3 | 6 | 7 | 11 | 3 |  |  |  |
| France | 17 | 2 | 2 | 6 | 4 |  | 1 | 3 | 1 | 1 |  | 1 |  |  |
| Spain | 12 | 5 |  | 3 | 3 | 4 | 1 | 2 | 3 | 1 |  |  |  |  |
| Albania | 8 | 0 | 2 | 1 |  |  |  | 2 | 3 |  |  |  |  |  |
| Algeria | 4 | 0 |  | 1 | 1 | 1 |  |  |  | 1 |  |  |  |  |
| Croatia | 4 | 0 | 2 |  |  |  |  |  | 1 | 1 |  |  |  |  |
| Cyprus | 3 | 0 | 1 | 1 |  |  |  |  | 1 |  |  |  |  |  |
| Montenegro | 2 | 0 | 1 | 1 |  |  |  |  |  |  |  |  |  |  |
| Syria | 2 | 0 |  |  |  |  |  |  | 2 |  |  |  |  |  |
| Tunisia | 2 | 0 |  |  |  |  |  |  |  | 2 |  |  |  |  |
| Egypt | 1 | 0 |  |  |  |  |  |  | 1 |  |  |  |  |  |
| **Totals** | 207 | 29 | 28 | 36 | 24 | 8 | 9 | 39 | 42 | 37 | 7 | 3 | 0 | 3 |

Forty five records contained both length and weight, which allowed us to fit length-weight relationships (Figure 3C) and predict these measurements when either one was missing. With these models, we predicted individual length for 17 additional records and weight for 36 records for a total of 98 individuals. Using these data, we were able to estimate individual ages with the von Bertalanffy growth curve (Figure 3B). Both size and age distributions showed that juveniles were caught more often than adults.

**Figure 3.**
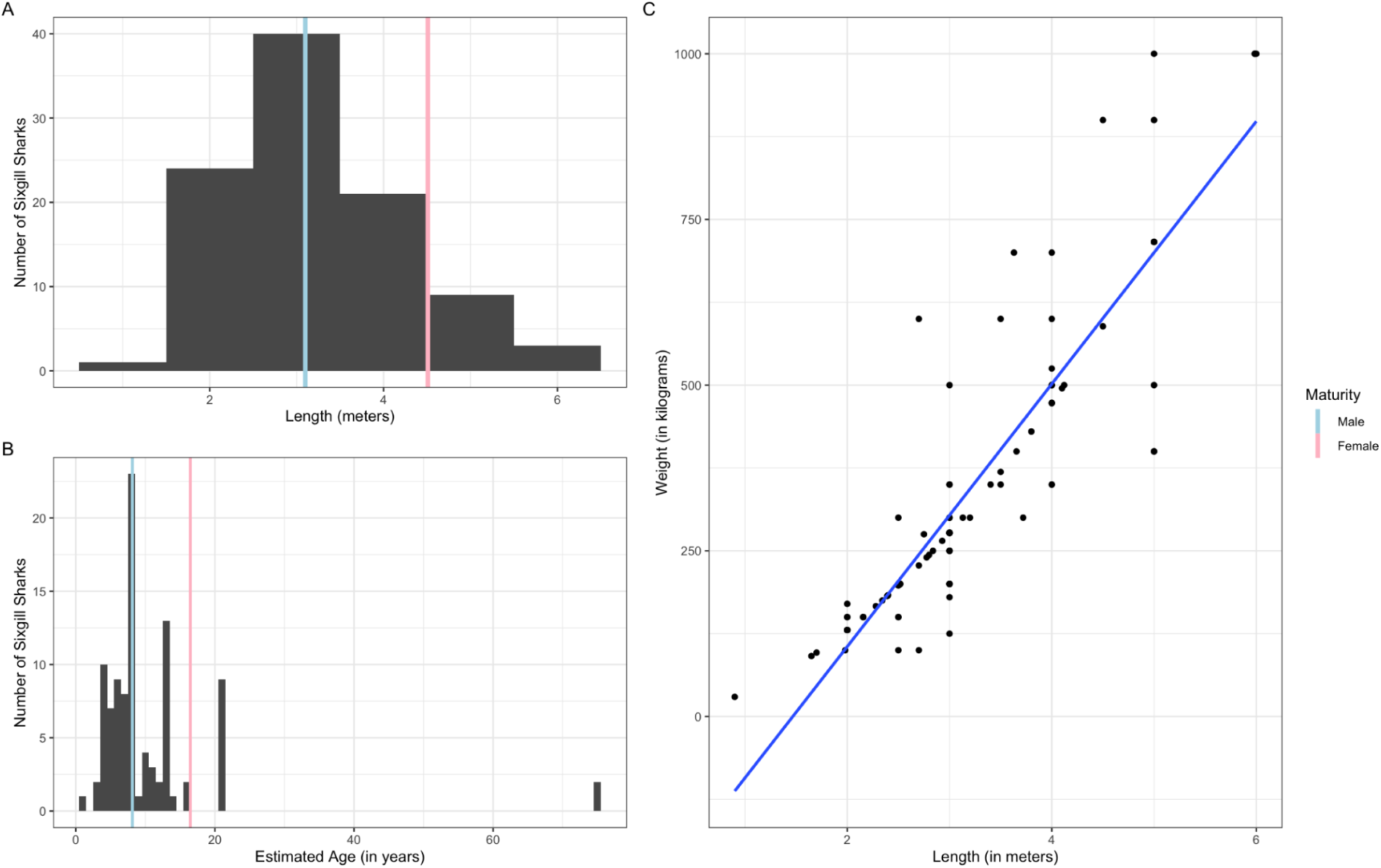
(A) Histogram of 98 *H. griseus* entries from sharkPulse that have total length either reported or estimated from length-weight relationship; (B) Histogram of ages of the 98 entries estimated from the von Bertalanffy growth function using FishBase coefficients; (C) Graph of length-weight relationship that provided estimates of missing lengths and weights where a = 6.90932 and b = 1.858319818

The factorial GLM yielded significant results for latitude and longitude as well as for years 2016 through 2021 in comparison to the intercept, year 2012 (p < 0.05, Table 3). Additionally, longitude and year were considered highly significant in the continuous generalized linear model (p << 0.05, Table 3) with latitude being near significant (p = 0.0526, Table 3). By graphing these two models together, we see that the continuous GLM portrayed by the line and ribbon shows an overall decreasing population trend for *H. griseus* from 2012 to 2021. There was a 93.13% decrease in the predicted abundance of *H. griseus* within the past decade (Figure 5) with a prominent hotspot in the Aegean Sea from 2015-2017 and a one around Algeria in 2015 (Figure 6). The temporal model also predicted two small increases in the *H. griseus* population during 2013 and 2017 (Figure 5) with 2017 being the only statistically significant (p < 0.05, Table 3) year out of the two. The time series produced from spatially mapping the GAM indicates a downward trend as well with spatial variability, namely in regions around Spain, Italy, and Greece (Figure 6). There was a lack of data starting in 2019 due to the pandemic which is why there is not a continuous plot of distribution for 2020 and 2021.

**Table 3.** Summary statistics of the factorial GLM, continuous GLM, and factorial GAM results testing sixgill abundance as a factor of longitude, latitude, and year.

| Factorial GLM (AIC = 1005.1) |  |  |  |  |
| --- | --- | --- | --- | --- |
|  | Estimate | Std. Error | z | P |
| (Intercept) | -4.110236 | 1.076176 | -3.819 | 0.000134 *** |
| latitude | 0.056154 | 0.02432 | 2.309 | 0.020945 * |
| longitude | 0.042623 | 0.007825 | 5.447 | 5.12e-08 *** |
| year2013 | 0.204584 | 0.255239 | 0.802 | 0.422821 |
| year2014 | -0.074723 | 0.276203 | -0.271 | 0.786747 |
| year2015 | -0.70938 | 0.407588 | -1.74 | 0.081783 . |
| year2016 | -0.948716 | 0.388584 | -2.441 | 0.014628 * |
| year2017 | -0.542776 | 0.252082 | -2.153 | 0.031305 * |
| year2018 | -0.959545 | 0.249528 | -3.845 | 0.000120 *** |
| year2019 | -1.029584 | 0.254958 | -4.038 | 5.39e-05 *** |
| year2020 | -1.97891 | 0.425981 | -4.646 | 3.39e-06 *** |
| year2021 | -2.678107 | 0.488015 | -5.488 | 4.07e-08 *** |
| Continuous GLM (AIC = 1007.8) |  |  |  |  |
|  | Estimate | Std. Error | z | P |
| (Intercept) | 440.789987 | 46.469202 | 9.486 | < 2e-16 *** |
| latitude | 0.045298 | 0.023376 | 1.938 | 0.0526 . |
| longitude | 0.043355 | 0.007448 | 5.821 | 5.85e-09 *** |
| year | -0.220782 | 0.023009 | -9.596 | < 2e-16 *** |
| Factorial GAM (AIC = 930.5131) |  |  |  |  |
|  | Estimate | Std. Error | z | P |
| (Intercept) | -2.26084 | 0.29683 | -7.616 | 2.61e-14 *** |
| year2013 | 0.15726 | 0.26188 | 0.601 | 0.548157 |
| year2014 | -0.01095 | 0.2868 | -0.038 | 0.969549 |
| year2015 | -0.63072 | 0.43218 | -1.459 | 0.144458 |
| year2016 | -0.8327 | 0.39782 | -2.093 | 0.036335 * |
| year2017 | -0.39988 | 0.26373 | -1.516 | 0.129461 |
| year2018 | -0.65945 | 0.25598 | -2.576 | 0.009990 ** |
| year2019 | -0.90036 | 0.26975 | -3.338 | 0.000845 *** |
| year2020 | -1.61635 | 0.43676 | -3.701 | 0.000215 *** |
| year2021 | -2.27747 | 0.49805 | -4.573 | 4.81e-06 *** |
|  | edf | Ref.df | Chi.sq | P |
| s(latitude,longitude) | 28.27 | 28.87 | 84.11 | 5.15e-08*** |

The compiled historical data were published from 1883 to 2012 with most studies focusing on trawl and longline fisheries (Table 1). The data came from multiple different sources but mainly stemmed from fisheries surveys. There was an increasing trend of the perceived abundance of *H. griseus* in various regions of the Mediterranean starting from very rare and rare occurrences in the late 1800s and becoming common in the early 2000s. By quantifying the perceptions and averaging them, we were able to see a slow rise throughout the entire Mediterranean Sea over 50-year periods (Figure 4).

**Figure 4.**
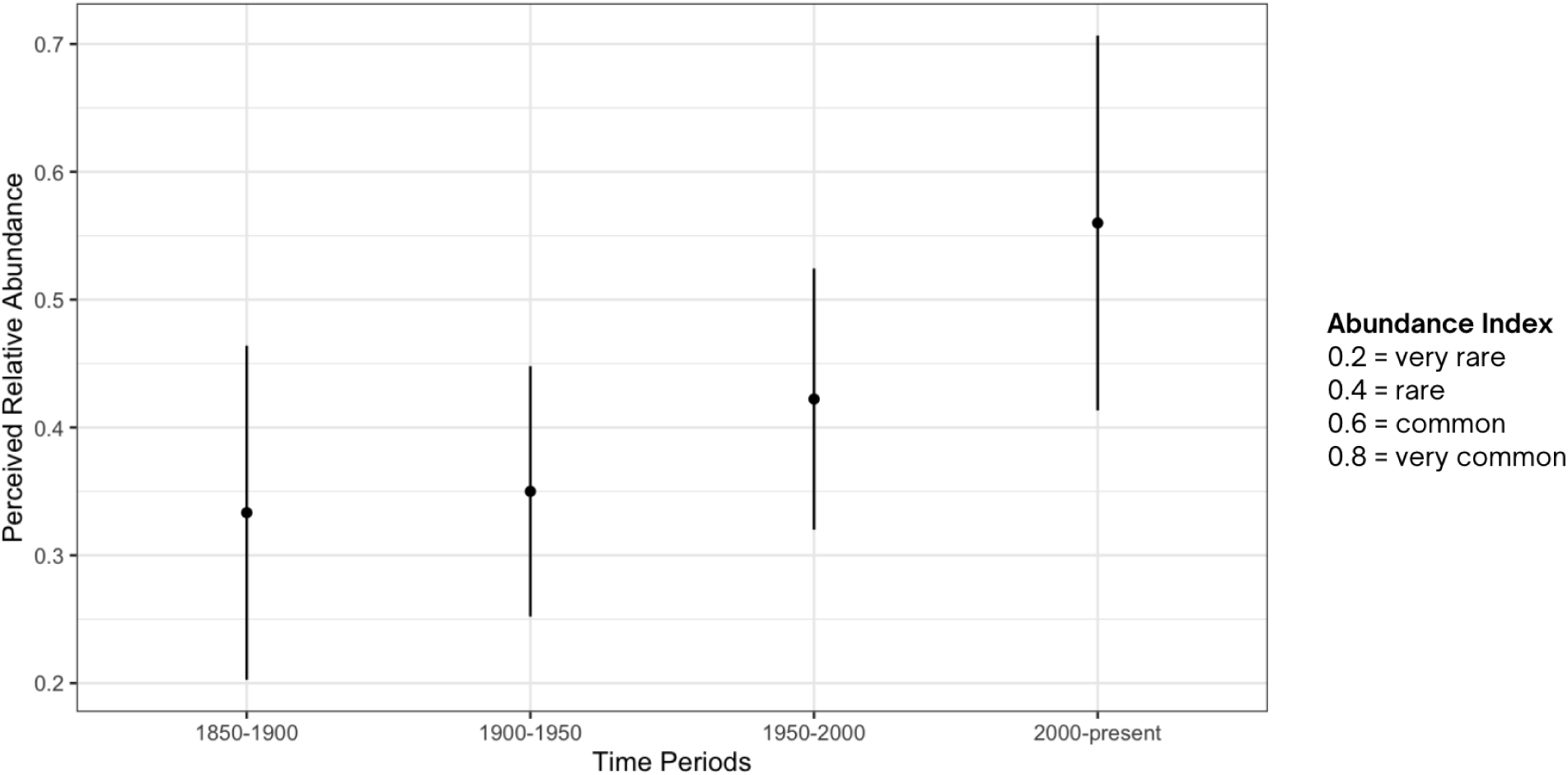
Temporal trend showing an increasing trend of perceived abundance based on historical literature review. The mean and standard error of each 50-year grouping was calculated based on Table 1’s four-level class coding.

## DISCUSSION

Fishing exploitation has a strong direct and indirect effect on exploited populations, bycatch species and their ecosystems. In the Mediterranean Sea, humans have severely depleted virtually all large predatory sharks once occurring in the region, preluding important ecosystem consequences. Here we tested the effect of these changes by analyzing an indicator species for these ecosystem effects. Despite the IUCN global assessment categorizing *H. griseus* as Near Threatened (Finucci *et al*. 2020), the Mediterranean regional assessment lists the species as Least Concern with a stable population which contradicts the centuries-old overexploitation of marine resources that has occurred in the region. While we were not able to reconstruct an entire time frame of population trends with the same type of data given the unknown longevity of this species, we were able to piece together a quadratic trend on the abundance of *H. griseus* using qualitative historical literature and quantitative opportunistic data.

We detected a 93.13% decline in the population of *H. griseus* within the past decade (Figure 5) which is indicative of potentially increasing fishing pressure that threatens the current “safe” conservation status of the species. Similar to the different hot and cold spots in our spatial analysis (Figure 6), fishermen who were interviewed in Greece and northeastern Spain believed that the population of *H. griseus* was stable while those in Albania, Tunisia and Cyprus thought the abundance was decreasing (Nuez *et al*. 2023). Over 100 records of *H. griseus* dating from 1908 to 2022 were documented in the Adriatic Sea, making it a common species (Soldo and Lipej 2022), which was not seen in our spatial analysis. Specifically, in the western Mediterranean where trawling is the most important, fisheries have expanded from the continental shelf to the slope in an effort to capture higher economically-valued species (Ramirez-Amaro *et al.* 2020). This, in turn, increases the chances of *H. griseus* being landed as bycatch which is evident in the western portion of the Mediterranean Sea having consistently fewer individuals compared to the eastern half (Figure 6). Environmental factors such as depth, sea bottom temperatures and habitat preferences could have caused populations to congregate in certain areas (Ruiz-Garcia *et al.* 2023) while anthropogenic factors such as the unequal distribution of vessels in the Mediterranean Sea, targeted species, and choice of fishing gear (Fiorentino and Vitale 2021) could play a role in exacerbating vulnerable populations of *H. griseus*.

**Figure 5.**
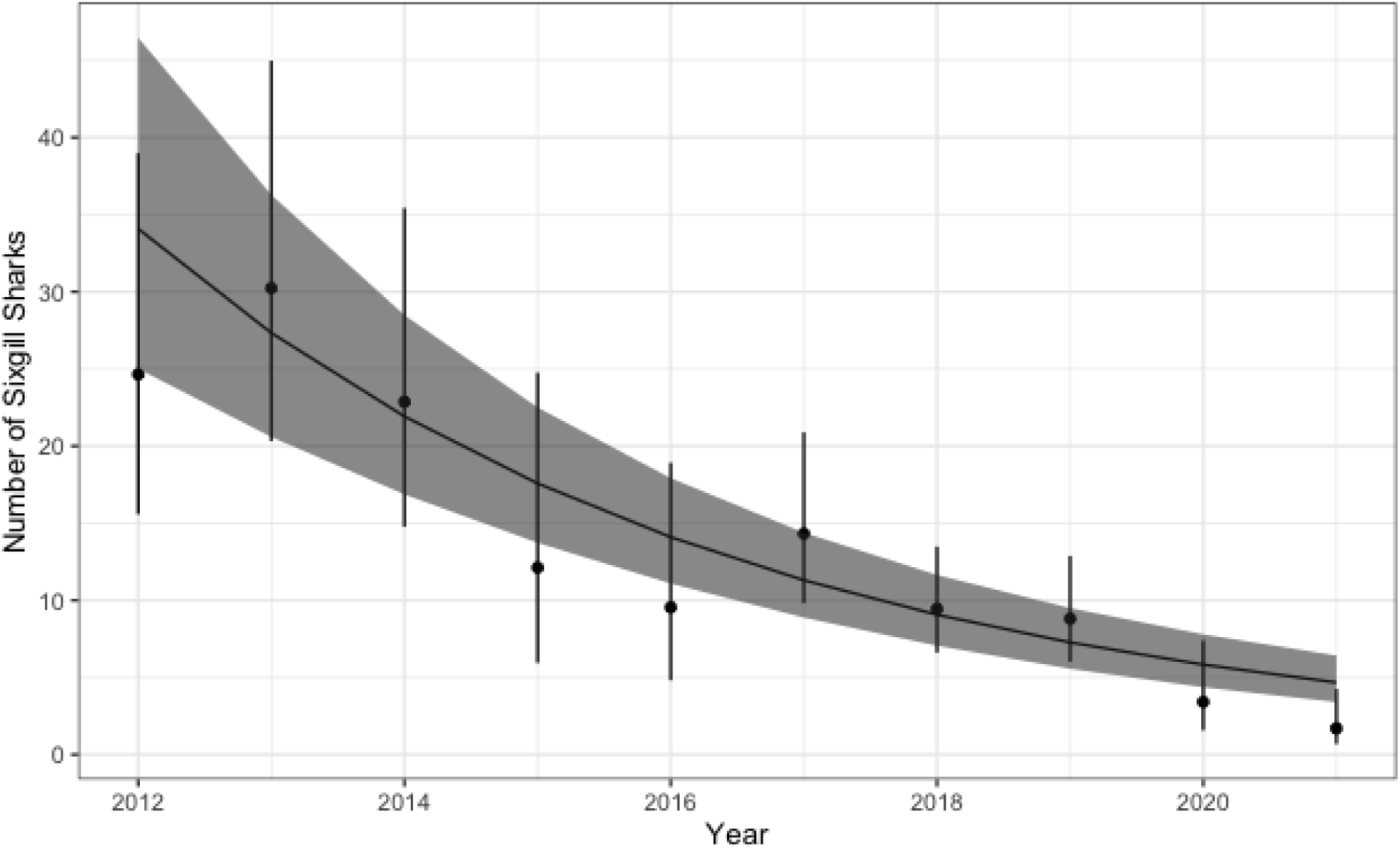
Temporal trend from 2012 to 2021 predicted from a GLM of *Hexanchus griseus* sightings as a function of latitude, longitude, and year with an observation effort of all other sharks in the Mediterranean. Trend was estimated from the parameter estimates of the GLM; latitude and longitude were fixed to plot these trends in a two dimensional plot

**Figure 6.**
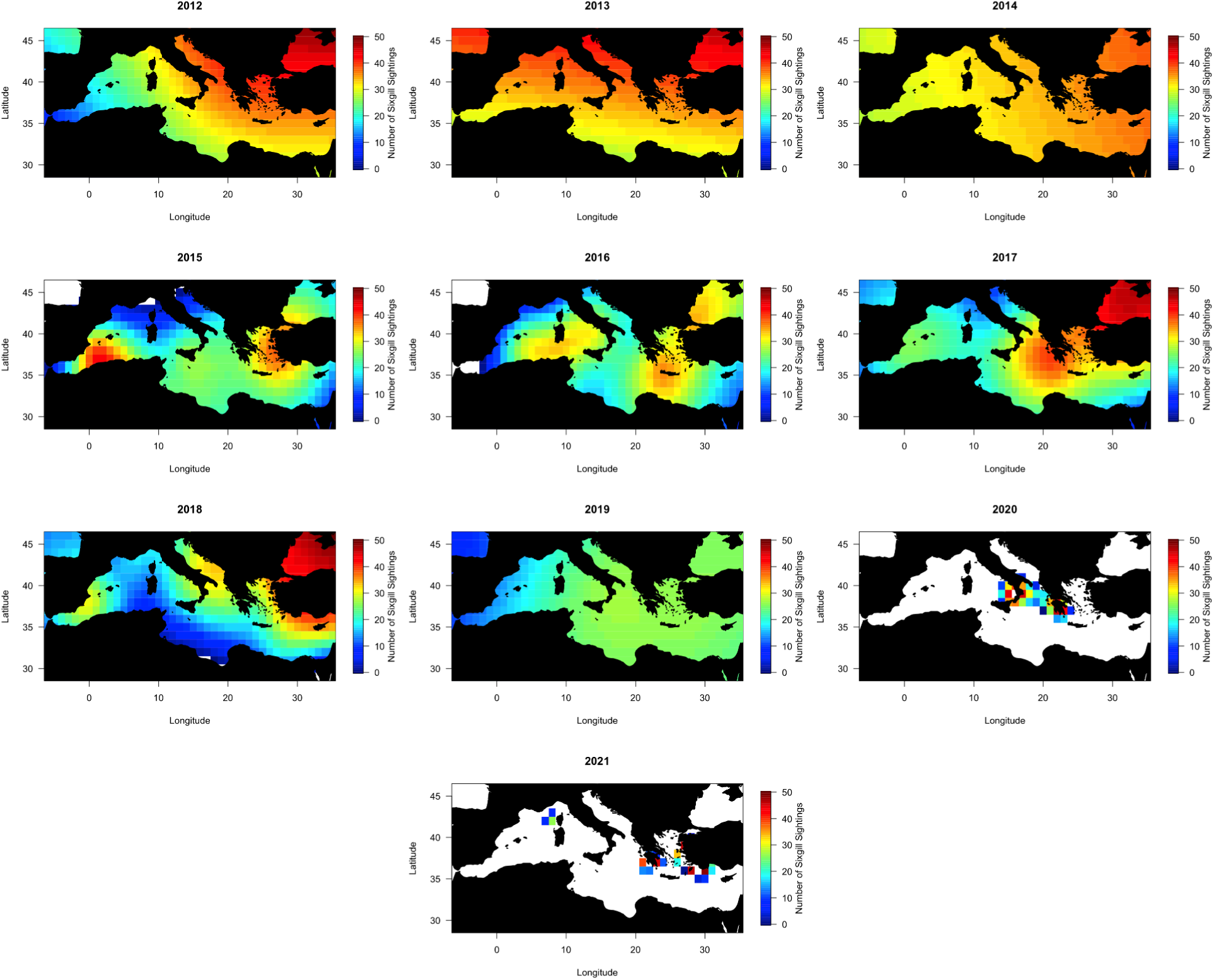
Time series of the predicted distribution of *H. griseus* based on the generalized linear model. The prominent white space in the 2020 and 2021 maps are due to lack of data from those years.

Although the most important age class to protect are juveniles nearing maturity (Simpfendorfer 1999), we found that a majority of the *H. griseus* individuals caught or observed were immature specimens (Figure 3) which can lead to undesirable consequences for the population. This has been seen in other studies surveying *H. griseus* both in and out of the Mediterranean Sea (Mili *et al.* 2021, Kabasakal 2013). Juveniles, despite preferring deep-water, were found to occupy a higher position in the water column (Griffing *et al.* 2019); some were even common in shallow coastal waters (Mili *et al.* 2021) because they were following their prey or avoiding predation and competition from adults that could be cannibalistic (Carey and Clark 1995). Their depth occupation in the water column results in a stronger likelihood of being captured by fishing gear that adults are more capable of evading or breaking through. On the other hand, fisheries could potentially be landing more juveniles because there are simply more of them in the water. Unfortunately, populations mainly composed of juveniles are considered strong evidence of depletion caused by overfishing and should warrant a closer examination of fishing impact. Although we were not able to determine a sex ratio from the sharkPulse records due to a very low sample size, many studies have found a sex ratio skewed towards females (Kabasakal 2004(1:2.25), 2006(1:2.54), 2013 (1:2.61); Capapé *et al.* 2003 (1:1.45); Celona *et al.* 2005 (1:1.69)). This could disproportionately affect which sex is landed more often by fisheries, leading to fewer viable females in the population. High juvenile mortality rates coupled with a biennial reproductive cycle and overfishing of adult populations can stall reproductive efforts, lower overall recruitment and eventually lead to entire populations dying off.

To compare the recent trends with baseline population levels, we collected historical information from various regions of the Mediterranean Sea to provide insights on periods not covered by the sharkPulse data and found that the qualitative measurements of *H. griseus* abundance were increasing (Figure 4). During the late 18th and throughout the 19th century, the practices surrounding fishing had begun to evolve in the countries surrounding the Mediterranean Sea. The continuous migration of communities and increasing demand from marketplaces all over the basin led to rapidly changing techniques. These changes combined with pollution from industrial revolution byproducts accelerated the exploitation of marine resources (Faget and Sacchi 2014). Despite this centuries-old overexploitation, the perception was that *H. griseus* was fairly rare up until the 2000s (Table 1). In multiple sectors like the Adriatic and Tyrrhenian Sea, fishermen who were involved in trawl and/or longline fisheries scarcely saw *H. griseus*. Concurrently, other large predatory sharks dramatically decreased in abundance over the past two centuries with some species showing rates of decline greater than 96% (Ferretti *et al.* 2008). This could have provided *H. griseus* an opportunity to rise from the deep region in the water column they originally occupied. We see this shift occurring in the historical data as *H. griseus* becomes more common starting from a study period that began in the 1980s (Hadjichristophorou 2006) which focused on the north and west coasts of Cyprus (Table 1). The Mediterranean Sea has a very distinct longitudinal biodiversity gradient which decreases from west to east (Coll 2010), which could explain why *H. griseus* was first more observed in the east. The lack of competition and predation from a less biodiverse region could have allowed *H. griseus* to expand in its range. It is also important to note that depending on the region, *H. griseus* had extremely variable listings in regards to its value. For example, it was listed as non-commercially important in Cyprus (Hadjichristophorou 2006) but in Syria, it was very important economically and highly consumable (Saad *et al.* 2006). The difference in value can lead to the species being merged with other cartilaginous fishes during reports (Schembri 2006) which can be detrimental when attempting to study their abundance.

A species like *H. griseus* is very difficult to study *in situ* due to its wide depth range and distribution. By using sharkPulse, we were able to extract information from opportunistic data over a long period of time. We were able to predict changes occurring over the vast region of the entire Mediterranean Sea which is very important given the spatial variability of *H. griseus* throughout the past two centuries. Each country that borders the Mediterranean Sea had their own perceptions of the abundance of *H. griseus* both in the past and present, which is understandable given the different fishing gear(s) used and the underwater habitats available in a certain area. Although this study focused on the Mediterranean Sea, future studies can use the same or similar opportunistic data in many different areas around the world. It is also important to take into account historical data since many marine ecology papers tend to focus too narrowly on recent years (Fortibuoni *et al.* 2010). By integrating historical data with modern quantitative research, we can drastically increase our knowledge of current and past populations of various marine organisms.

In conclusion, this study aimed to describe the population trends of *H. griseus* in the Mediterranean by taking advantage of opportunistic data and supplementing the data we recovered with historical research about the species. By using a crowd-sourcing platform like sharkPulse, we were able to extract information not only about the abundance and distribution of the species but also information about their morphological traits like their size, weight, sex, age, etc. Using these kinds of population trends could benefit conservation efforts and inform decisions that need to be made in regards to the unchecked fishing pressures present in the Mediterranean Sea. Despite the increasing trend of *H. griseus*’s perceived abundance in the 1800s and 1900s, more modern times have shown a steady decrease in their abundance. This decrease is most likely due to the increasing fishing pressures that are fueled by an ever-growing human population. Because of *H. griseus*’s Near Threatened/Least Concern (regionally) categorization by the IUCN (Finucci *et al*. 2020, Soldo *et al.* 2016), research efforts may not be strictly focused on this species. However, if we neglect *H. griseus* and other species in similar situations, we could miss the chance to protect and conserve them and they could be gone in the blink of an eye. As climate change and subsequent consequences become increasingly problematic, the overexploited Mediterranean Sea will have even higher levels of extinction risk compared to the already-elevated level in 1980 (Walls and Dulvy 2021) which could deeply hurt organisms with slow life history traits like sharks.

## ACKNOWLEDGEMENTS

This research would not have been possible without the help of the SeaQL Lab headed by Francesco Ferretti and the sharkPulse monitoring platform created by the lab. I would also like to thank Jeremy Jenrette and Chiara Gambardella for their helpful comments and suggestions on the manuscript and for their everlasting support in general.

## Author contributions

ILJ and FF conceived the study. ILJ conducted data mining, analyses, and wrote the manuscript. FF supervised and contributed to data interpretation, analysis, and writing.

